# Optimization of a deep mutational scanning workflow to improve quantification of mutation effects on protein-protein interactions

**DOI:** 10.1101/2023.10.23.563542

**Authors:** Alexandra M. Bendel, Kristjana Skendo, Dominique Klein, Kenji Schimada, Kotryna Kauneckaite-Griguole, Guillaume Diss

## Abstract

Deep Mutational Scanning (DMS) assays are powerful tools to study sequence-function relationships by measuring the effects of thousands of sequence variants on protein function. During a DMS experiment, several technical artefacts might distort non-linearly the functional score obtained, potentially biasing the interpretation of the results. We therefore tested several technical parameters in the deepPCA workflow, a DMS assay for protein-protein interactions, in order to identify technical sources of non-linearities. We found that parameters common to many DMS assays such as amount of transformed DNA, timepoint of harvest and library composition can cause non-linearities in the data. Designing experiments in a way to minimize these non-linear effects will improve the quantification and interpretation of mutation effects.

## Background

How information encoded in DNA sequence is translated into molecular function is central to our understanding of the genotype-phenotype relationship. Deep Mutational Scanning (DMS) assays allow to evaluate the effect of thousands of sequence variants on a given molecular function in parallel and have therefore become powerful tools to address this question. In DMS assays, a variant library is created by mutagenizing a sequence of interest and exposing this library to a selective assay that separates variants based on their activity. Because the assay is performed in a pooled format, high-activity variants are enriched under the selective conditions while low-activity variants are depleted. Enrichment and depletion are quantified using deep sequencing [1,2].

Proteins perform most biochemical functions within a cell and do so by interacting with one another. The specificity of these protein-protein interactions (PPIs), i.e. the information about which two proteins interact with one another, is encoded in the DNA sequence and is therefore central to the sequence-function relationship. Moreover, it has been shown that disease-associated variants are enriched at interaction interfaces, which is why perturbations of PPIs are of major interest in medical and pharmacological research [3].

The fast-progressing advances made in sequencing technology enable the discovery of more and more variants in the human population with potential effects on PPI affinity. The vast amount has long surpassed a number feasible to study individually. Recent publications have shown that DMS assays are very well suited for the study of PPIs or protein-ligand interactions at large scale [4–10].

deepPCA is one DMS approach to measure PPIs that proved to be very useful to study determinants of specificity in conserved protein interaction domains such as human basic leucine zippers [5,11]. In this assay, a Dihydrofolate Reductase-based Protein-fragment Complementation Assay (DHFR-PCA) is combined with DMS by performing DHFR-PCA in a pooled format and measuring the effect of thousands of mutations in a protein of interest on its ability to interact with its partners. DHFR-PCA is based on the reconstitution of a methotrexate (MTX)-insensitive mouse DHFR variant (Figure 1A). Both halves of the enzyme (referred to as DH-tag for the N-terminal fragment and FR-tag for the C-terminal fragment) are used to tag each of the two proteins of interest and expressed in *Saccharomyces cerevisiae* cells. Interaction between these two proteins promotes complementation between the two DHFR fragments and sustains growth in presence of MTX, which inhibits the endogenous DHFR [12,13].

**Figure 1.**
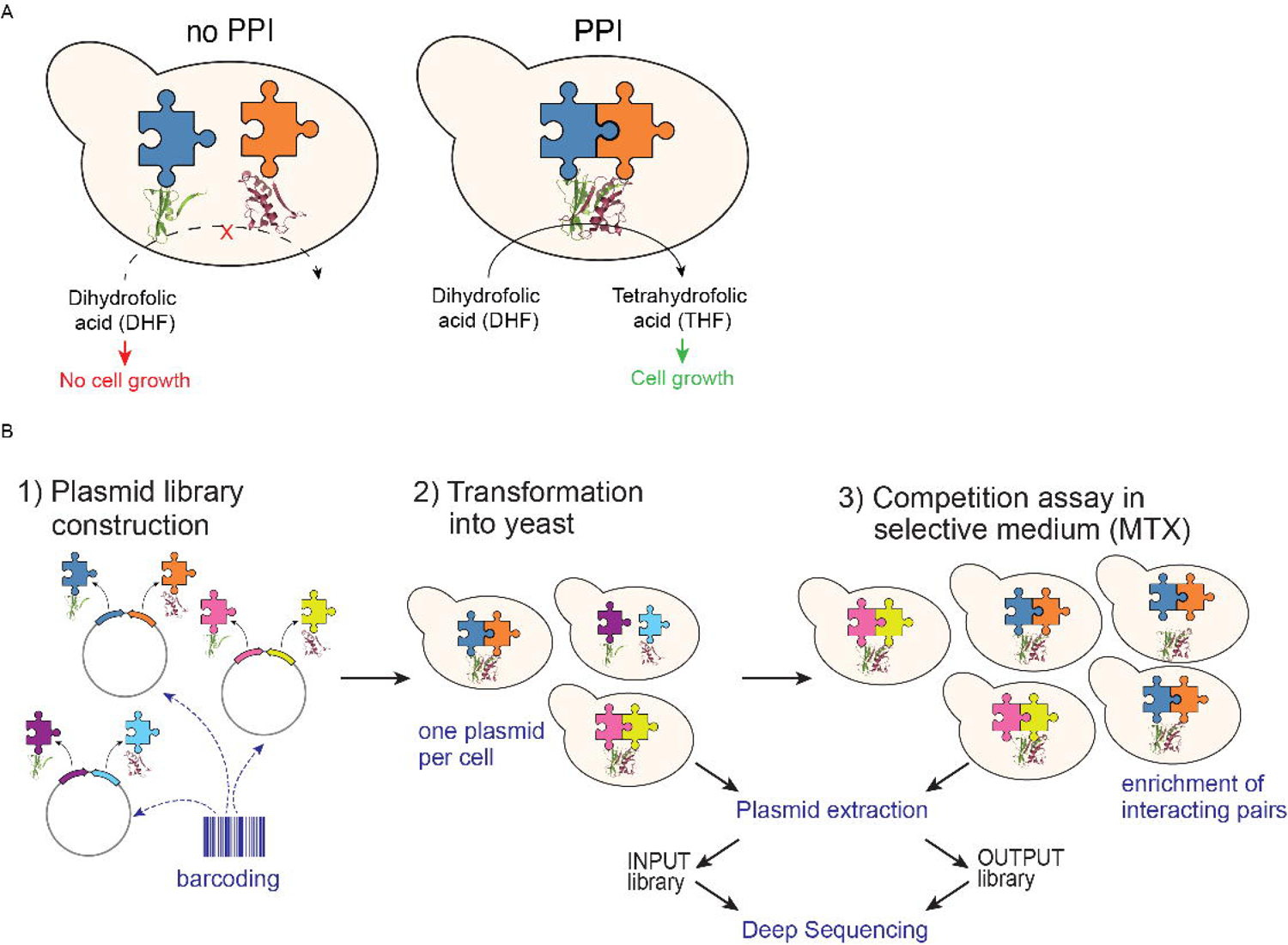
The deepPCA general principle and workflow. A. In DHFR-PCA, proteins of interest are tagged with the two opposite halves of a MTX-insensitive DHFR variant. Upon interaction of the tagged proteins, DHFR is reconstituted and functional. Yeast cells expressing the interacting pair can sustain cell growth in presence of MTX [13]. B. deepPCA: pairs of proteins tagged with opposite DHFR termini are expressed from single plasmids. Molecular barcodes encoded on the plasmids allow identification of the protein pair. Plasmids are transformed into yeast cells so that each cell expresses one plasmid. Cells expressing interacting proteins are enriched in selective medium containing MTX. Enrichment and depletion of interacting and non-interacting pairs is quantified by barcode sequencing of plasmids extracted from Input and Output cultures[11].

DHFR-PCA is quantitative, i.e. the growth rate of the respective cells depends on the concentration of reconstituted DHFR, which is itself a function of protein abundance and the affinity of the interaction [14]. In deepPCA, this quantitative aspect is leveraged in a pooled setting, where variants of the interacting proteins are enriched or depleted depending on the strength of their PPI.

deepPCA has been designed to allow library-on-library screening. First, intermediate libraries for DH- and FR-fusions are cloned with random DNA barcodes. The association between barcode and variant is determined for each library by deep sequencing and the two libraries are then combined on the same plasmid in a way that juxtaposes the barcodes. This plasmid library is then transformed into yeast cells at a low multiplicity of transformation, i.e., to minimize the number of cells with two or more plasmids. Competitive growth in MTX-containing medium then results in enrichment of interacting pairs and depletion of non-interactors. Input and output cultures (before and after competition) are harvested, and plasmids are extracted. Enrichment and depletion of individual pairs are quantified by deep sequencing of the molecular barcodes encoded on the plasmids. Frequencies of each pair in input and output populations are determined and used to calculate the growth rate (generations per hour) of cells expressing each pair of proteins as a proxy for the concentration of the complex (Figure 1B).

deepPCA’s readout, like many other DMS assays, maps non-linearly to the latent trait where mutations are expected to have additive effects [5,15]. Indeed, the readout is directly proportional to the concentration of the complementation complex, which is itself non-linearly related to the underlying energetic dimension where mutations that are not energetically coupled affect the free energy of the complex additively. Mutations that have additive effects at the energetic level will thus show non-additive effects at the level of protein complex concentration, which can be mis-interpreted as the two mutations having some structural or functional relationship. This phenomenon has been referred to as global, phantom or non-specific epistasis [5,16–18]. Regressing out these non-linearities is therefore required to access mutational effects on the latent additive trait. This is typically achieved by employing an appropriate mechanistic model describing the system, such as two- or three-states thermodynamic models for PPIs [5,6,11,17,19].

The experimental pipeline can also introduce additional non-linearities when technical factors affect different variants in different ways depending on their position within the dynamic range of the assay. These non-linearities can sometimes be accounted for even when not explicitly included in the thermodynamic model, for instance by being incorporated in some global parameters determining the shape of other non-linearities. However, they can still have negative effects by restricting the dynamic range of the assay. It is therefore important to identify potential sources of non-linearity in the workflow to improve the quantification of mutational effects.

The purpose of this study is thus to perform optimization experiments at different critical steps of the experimental pipeline to identify sources of non-linearity in the deepPCA workflow. The parameters tested were 1) amount of transformed library DNA, 2) MTX concentration, 3) timepoint of harvest, and 4) library composition. This improved our understanding of both the biological/physical and technical non-linearities and will allow us to account and correct for these sources of error in our experimental design. The improved data quality will enable a more accurate thermodynamic modelling of the underlying biophysical processes, benefiting a deeper understanding of evolutionary, epistatic, and pathogenic effects of mutations.

## Results and Discussion

### A yeast transformation protocol with minimal double transformants and optimized library coverage

If a cell contains two plasmids expressing proteins with different interaction strengths, the cell growth rate is mostly determined by the strong interaction pair and the change in frequency of the weak pair will be overestimated, which can be a source of non-linearity. Therefore, to preserve the quantitative nature of DHFR-PCA, the fraction of cells carrying two or more different plasmids must be minimized.

To determine the proportion of cells co-transformed by more than one plasmid in our large scale, high-efficiency transformation protocol, we transformed different amounts (between 250 ng and 1 μg) of an equimolar mixture of two centromeric plasmids (pRS415 and pdL00266) carrying each a different auxotrophic selection marker. We then plated on selective plates missing one or both corresponding metabolites to count the total number of transformed cells and the number of double transformants and calculated the percentage of cells carrying more than one plasmid (see Methods). Transformants are selected in two consecutive rounds for many generations before the start of the competition in the MTX medium, in order to overgrow the large majority of non-transformed cells. This should allow for cells carrying two different plasmids to lose one and thus decrease the percentage of double transformants [20]. To measure the rate of loss, we compared the percentage of cells carrying more than one plasmid directly after transformation and at the end of the selection process, i.e., just before inoculation of the MTX medium (Supplementary Table 1).

The percentage of double transformants directly after transformation and before MTX selection increased in a linear fashion with increasing DNA amount (Figure 2A). The total number of transformants also increases in a linear fashion when transforming with higher amounts of DNA (Supplementary Figure 1A). Moreover, it remained stable indicating that the cells maintain both plasmids over the multiple generations of growth.

**Figure 2.**
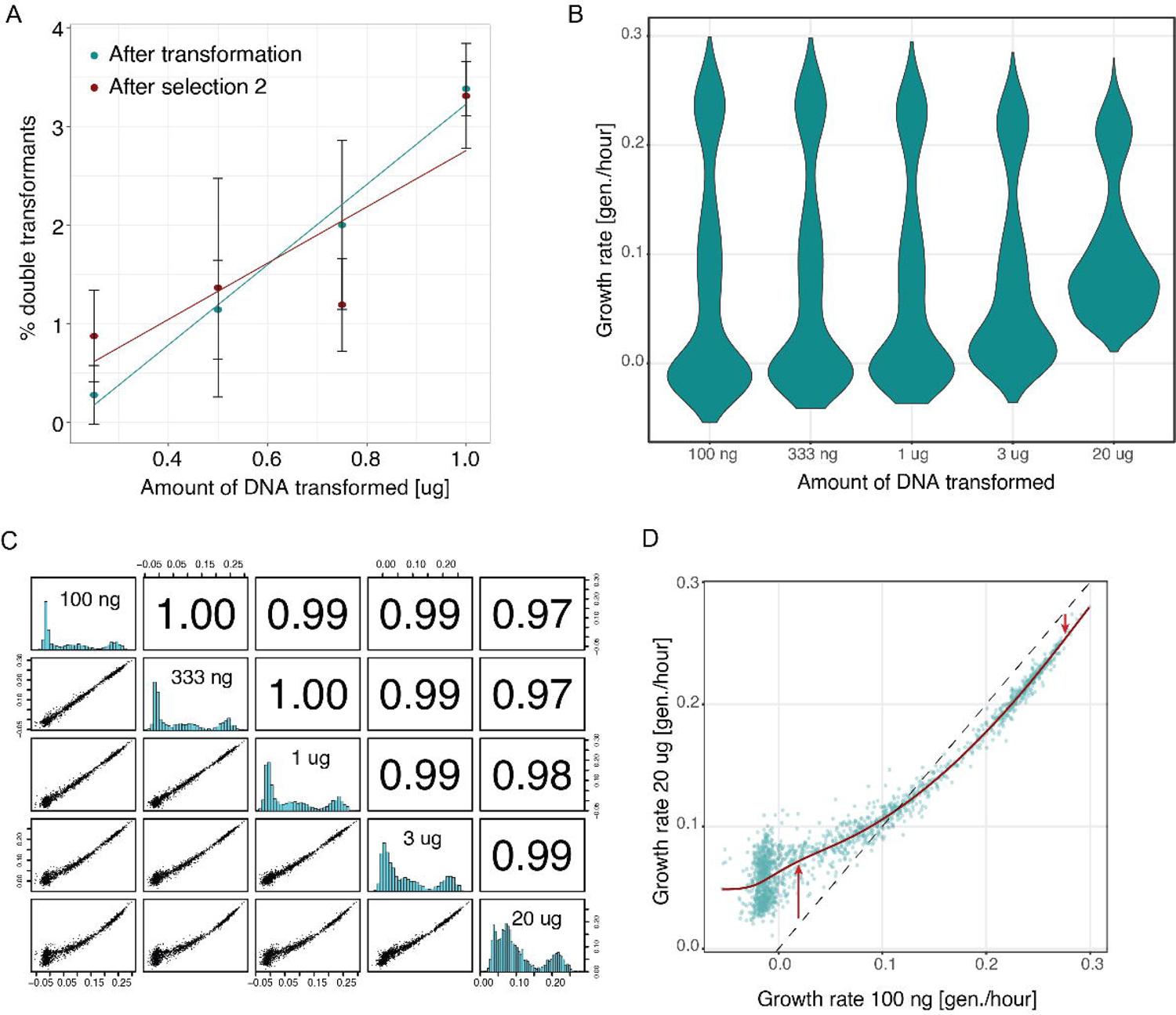
Effect of transforming with different amounts of DNA. A. Fraction of double transformants per cell for different amounts of DNA transformed. Cells were transformed with DNA amounts from 250 ng to 1 μg. The amount of double transformants was quantified after initial transformant selection and after selection round 2 (see Methods). B. Distributions of growth rates obtained from deepPCAs performed with cells transformed with differing amounts of DNA between 100 ng and 20 μg. C. Pairwise correlations of growth rates between samples from deepPCAs performed with cells transformed with differing amounts of DNA. Scatter plots are shown in the bottom left panels, histograms are shown on the diagonal, and Pearson correlation coefficients are shown in the upper right panels. D. Scatterplot between samples transformed with 100 ng DNA and 20 μg DNA.

We next measured the effects of transforming different DNA amounts and hence different percentages of double transformants on deepPCA interaction scores. We constructed a small library based on a JUN variant library recently assayed [11]. We combined the 608 single amino acid substitutions of the leucine zipper domain of human JUN (FR-tagged) with three out of 54 wild-type human basic leucine zippers (bZIPs; DH-tagged). The three wildtype bZIPs were selected to span a range of interaction strengths between JUN and its wild-type partners. FOS was used as a strong interactor, ATF7 as a weak interactor and NFE2 as a non-interactor. This library was used in most experiments presented in this paper and will be referred to as the balanced JUN library.

We performed deepPCA after transforming yeast cells with five different amounts of library DNA, expanding the range used above to better detect effects on interaction scores (100 ng, 333 ng, 1 μg, 3 μg, and 20 μg). We first compared the correlations between variant frequencies in input and output samples across all conditions. Overall, the correlations were very similar, but it was apparent that the output sample from the cells transformed with 20 μg of library DNA correlated stronger with all input samples than the others quantity transformed, indicating that library variants are less enriched or depleted when transformed with this high DNA amount (Supplementary Figure 1B; Supplementary Table 2). While input and output samples of all conditions clustered together rather randomly because of the high correlation, output samples from the 20 μg sample clustered together and separate from the other samples (Supplementary Figure 1B). To further investigate this, we calculated the per variant growth rate (generations per hour) for each sample and observed a narrower distribution in the 20 μg sample while the distribution in the remaining samples were very similar (Figure 2B; Supplementary Table 3). When looking at the correlations between growth rates, the correlation coefficients were very high but decreasing with increasing difference in DNA amount (Figure 2C). This is once more especially apparent for the 20 μg sample for which we observe a clear non-linear trend with growth rates of other samples. This can be explained by the expected high average number of plasmids per cell in that sample. For example, when directly comparing growth rates of the sample transformed with 100 ng DNA and the one transformed with 20 μg DNA, slow growers are most severely affected and grow faster in the high DNA sample (Figure 2D). This is because slow growers will likely get co-transformed with a plasmid that expresses a pair that interacts more strongly, and thus increase the growth rate of the cell in MTX beyond the level expected for the weakly interacting variant alone. There is only a slight decrease in growth rates for the strong interactors, which might be due to interference from the weaker co-transformed plasmid on the expression from the stronger plasmid. Importantly, these effects are not stochastic and affect the trend, as further demonstrated by the high correlation between output replicates of the 20 μg sample (Supplementary Figure 1C). Because of the large number of transformant per variant pair obtained (∼1000) and the high co-transformation percentage expected with 20 μg of DNA, each variant pair is expected to be co-transformed with a large fraction of all other variant pairs. The average effect thus becomes deterministic. On the contrary, if the average number of transformants per variant was low, a variant pair in one replicate might be co-transformed with strong variant pairs while in another replicate it might be co-transformed with weak variant pairs.

The non-linear trend and the narrowing of the distribution of interaction scores is still slightly apparent at 3 μg and negligible at 1 μg and lower amounts. We thus choose 1 μg as an optimal trade-off between the high number of total transformants required when working with large libraries and a low percentage of double transformants with minimal effects on the distribution of interaction scores.

### Increasing concentration of MTX results in higher but linear selection strength

Next, we tested the effect of MTX concentration during competitive growth. Previous deepPCA studies used a standard concentration of 200 μg/mL MTX [5,6,11][5,6], which we here refer to as 1X MTX. This concentration was first determined for performing DHFR-PCA on agar plates [13,21], and we therefore decided to verify that it is also well-suited for deepPCA in liquid culture. We evaluated the effect of 1/20X, ¼X, ½X, 1X, and 2X MTX concentration and a DMSO only control, controlling for the effects that some variants can have on actual cellular fitness, for instance by interfering with other cellular components. In addition, we also compared the results from different MTX vendors (using 1X MTX in the assay; Supplementary Tables 4 & 5).

We did not observe any effect from the supplier (Supplementary Figure 2A). We also did not observe any fitness effect in DMSO (Pearson correlation coefficient of 1.00 between input counts and output counts from cells grown in DMSO; Figure 3A, upper left). On the contrary, in presence of as low as 1/20X MTX, enrichments of some variants can be observed which is reflected in the lower correlation coefficients to input counts (Figure 3A). The correlation coefficients decrease together with increasing MTX concentration, indicating stronger selection at higher MTX concentrations. The correlation between output counts of the ¼, ½ 1 and 2X samples is 0.99, further indicating little differences in enrichments at different MTX concentrations.

**Figure 3.**
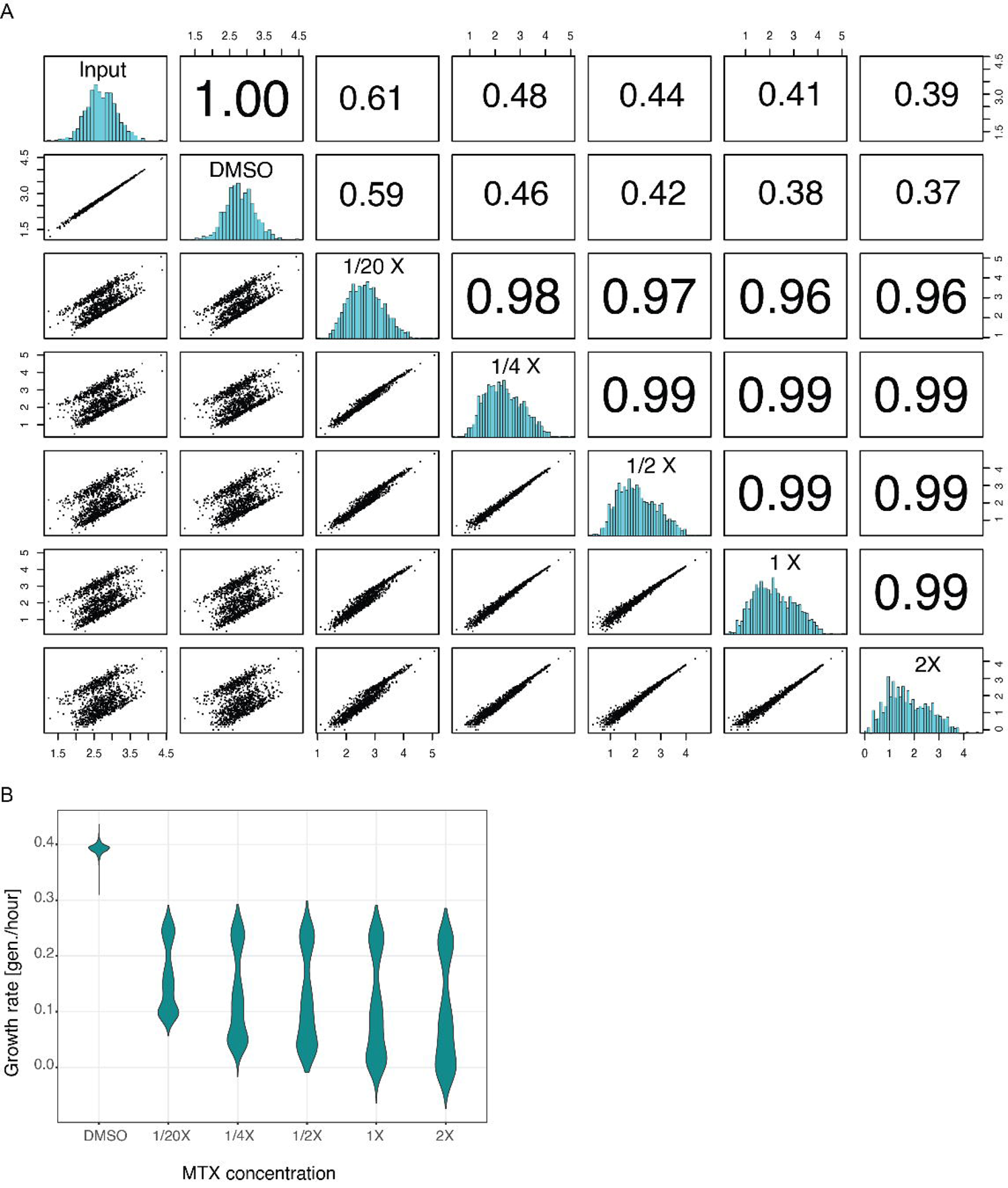
Different MTX concentrations do not result in non-linearities, but selection pressure is higher in higher concentration. A. Pairwise correlations of average counts between Input and Output samples of deepPCAs selected in DMSO and different concentrations of MTX. Scatter plots are shown in the bottom left panels, histograms are shown on the diagonal, and Pearson correlation coefficients are shown in the upper right panels. B. Distributions of growth rates of deepPCAs selected in DMSO and different concentrations of MTX.

Increasing MTX concentration stretches the growth rate distributions (Figure 3B), further indicating stronger selection regimes. The lower mode of the distribution decreases while the fast-growing variants are little affected, indicating a higher background growth at lower MTX concentration, probably resulting from an incomplete inhibition of the endogenous DHFR. Since the growth rates are perfectly linearly related at different MTX concentrations (Supplementary Figure 2B), increasing it from 1 to 2X does not confer significant benefit and would not justify the higher cost. However, decreasing MTX concentration 2X or 4X would have the advantage of a lower cost and might therefore be considered for very large and costly experiments. The slightly higher background growth can be accounted for by the global parameters of the thermodynamic model that linearly rescales the growth rate to the fraction bound [6]. Our results also show that a slight decrease in MTX concentration over the course of the experiment due to potential degradation would not affect growth rates non-linearly. We thus conclude that 1X is an optimal trade-off between cost and selection strength.

### The lag-phase at the start of the competition non-linearly affects growth rates

We next set out to determine the ideal harvesting strategy to maximize the size and spread of the measured effects, while keeping the error small. To this end, different harvesting time points for the input and output samples were considered. Previously, the input cultures were harvested before inoculating the MTX-containing selection medium. A portion of these cells was used to inoculate the competition culture which was grown for five more generations before harvesting the output sample. At the beginning of the growth in the MTX medium, cells have enough stores of tetrahydrofolate (THF), the product of DHFR, and downstream metabolites to sustain growth until relying solely on the complemented DHFR activity. Therefore, cells expressing different pairs of proteins are expected to grow at the same rate for a few hours. This is clearly apparent in screens on agar plates, where cells at all array positions were able to grow to form small colonies and only the positions at which cells expressed interacting proteins would grow further. This lag before the selection kicks in can thus potentially non-linearly affect the estimated growth rate such that the difference between true and estimated growth rate would be larger for weak interactors since the MTX-dependent growth time is overestimated. Strong interactors, on the other hand, might not see a strong decrease in growth rate upon MTX selection, and their estimated growth rate will be very close to the true growth rate (Figure 4A).

**Figure 4.**
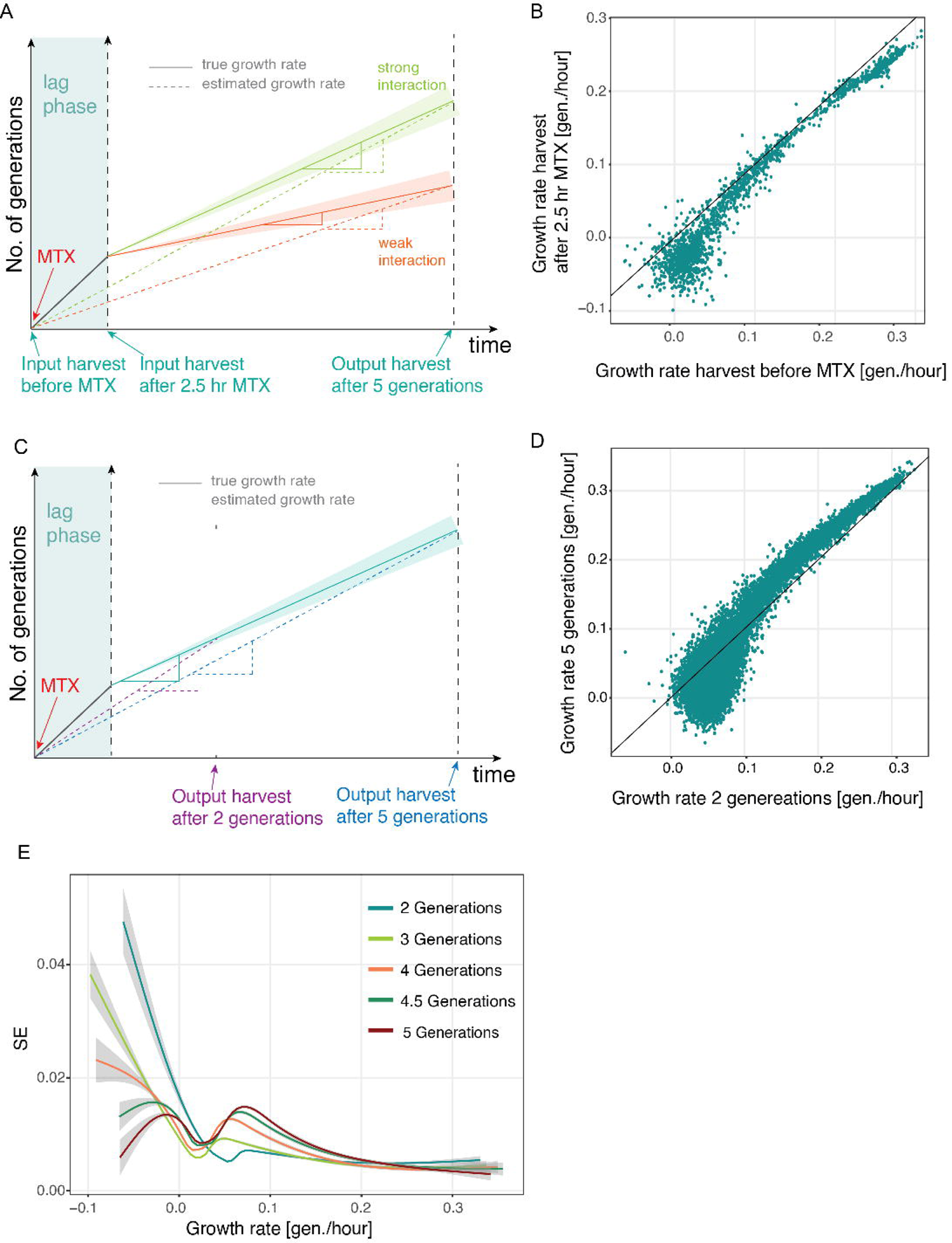
Growth time and pre-selection phase non-linearly affect growth rates. A. Harvesting Input samples before or after a pre-competition phase in which all yeast cells still grow at equal growth rates can result in differences in measured growth rate. Harvesting before pre-competition phase can result in an overestimation of growth rate of slow growers because the assumed duration of selection is longer than the actual duration. B. Scatter plot comparing growth rates of deepPCAs for which the Input sample has been harvested before or after the pre-competition phase (after 2.5 hours in MTX). C. Harvesting after shorter growth time can result in further overestimation of growth rate for slow growers because the pre-competition phase represents a larger fraction of the overall selection time and more strongly influences the estimated growth rate than after a long growth time. D. Scatter plot comparing samples of deepPCAs grown for 2 generations and 5 generations. E. Standard errors of growth rate as a function of growth rate for samples harvested after different generations.

To investigate this effect, we used the same JUN balanced library and compared growth rates obtained following the normal protocol to those obtained by first pre-incubated cells in the MTX medium for 2.5 hours before collecting the input sample and inoculating fresh MTX medium. Both output cultures were harvested at a cell density corresponding to a population growth of five generations. As expected, we do see a slight non-linear relationship between growth rates with a stronger effect on weak interactors, i.e., pairs with a lower growth rate that show a disproportionately stronger increase in generation time than the strong interactors (Figure 4B; Supplementary Tables 4 & 6).

Pre-incubation in MTX is therefore recommended to minimize non-linear effects. However, the extra steps make the experimental procedure more complex and costly and might add to the technical noise of the data while the non-linear effects can be accounted for by the global parameters of the thermodynamic model that control the curvature of the non-linear transformation[6]. The decision of whether or not to perform the pre-incubation is thus left to the user depending on the use of the data.

### The choice of harvesting timepoint is another source of non-linearities

The number of generations in competitive growth might also result in differences between estimated and true growth rate (Figure 4C). We therefore performed deepPCA by harvesting the input samples before MTX selection and harvesting the output samples after 2, 3, 4, 4.5 and 5 generations of total population growth (Supplementary Tables 7 & 8). For this experiment, we used the full JUN variant library combined with the 54 wt bZIPs from [11]. We first compared the distributions of generation times across the different samples. Though similar, the harvest after 2 generations shows a slight shift towards higher average growth rates indicating that the selective pressure was not applied long enough for strong depletion of the slow growers (Supplementary Figure 3A). Additionally, a non-linear relationship between the samples harvested after different generation times that seems to affect slow-growers disproportionately stronger is observed. The higher the difference in generations at harvest the more pronounced is this effect, being the strongest between the samples harvested at 2 and at 5 generations (Figure 4D, Supplementary Figure 3B). This can also be explained by the effect of the lag-phase, which is proportionately stronger when the effective competition is shorter (Figure 4C). For this reason, a harvest at later timepoint (i.e., after 5 generations) will result in estimated growth rates that are closer to the true growth rates than those estimated from samples harvested earlier.

On the other hand, with increasing time of growth, also the noise and therefore the error of measurement increases, especially for slow growers (Figure 4E). This is likely due to lower depletion of slow growers and thus higher read counts, the main source of measurement error[22,23]. Thus, choosing the timepoint of harvest is a tradeoff between how close to the true growth rates the estimated ones are and how high of an error and drop-out rate of slow growers is tolerable.

### Library composition

In principle, measured growth rates for individual protein pairs should be independent of library compositions. To test this, we performed deepPCA using our pre-determined standard settings of 1 μg of DNA for transformation, harvest of input samples before the competition and harvest of output samples after 5 generations of growth. We screened three different libraries of identical sizes, the balanced JUN library, a slow-growing library in which we replaced the strong interactor FOS by the weak interactor ATF2, and a fast-growing library in which we replaced the weak interactor NFE2 by the strongest interactor JDP2. Importantly, ATF7 was present in all three libraries, allowing to compare growth rates and test if they are affected by library composition (Supplementary Tables 9 & 10). Growth rates for the ATF7 pairs were well correlated between the different libraries (Figure 5A). However, their distributions differ, and the growth rates estimated for ATF7 pairs when competing against stronger pairs are lower (Figure 5B). This is also true for other partners that overlap only in two libraries. For example, FOS shows overall higher growth rates in the balanced library where it forms the strongest interactions with JUN variants than compared to the fast-growing library where JDP2 forms the strongest interactions with JUN variants. This is more likely to be due to technical artefacts than actual differences in growth rates, such as inaccuracies in the estimation of frequencies from read counts. However, since growth rates are linearly related across the different libraries, this is easily accounted for in the thermodynamic model.

**Figure 5.**
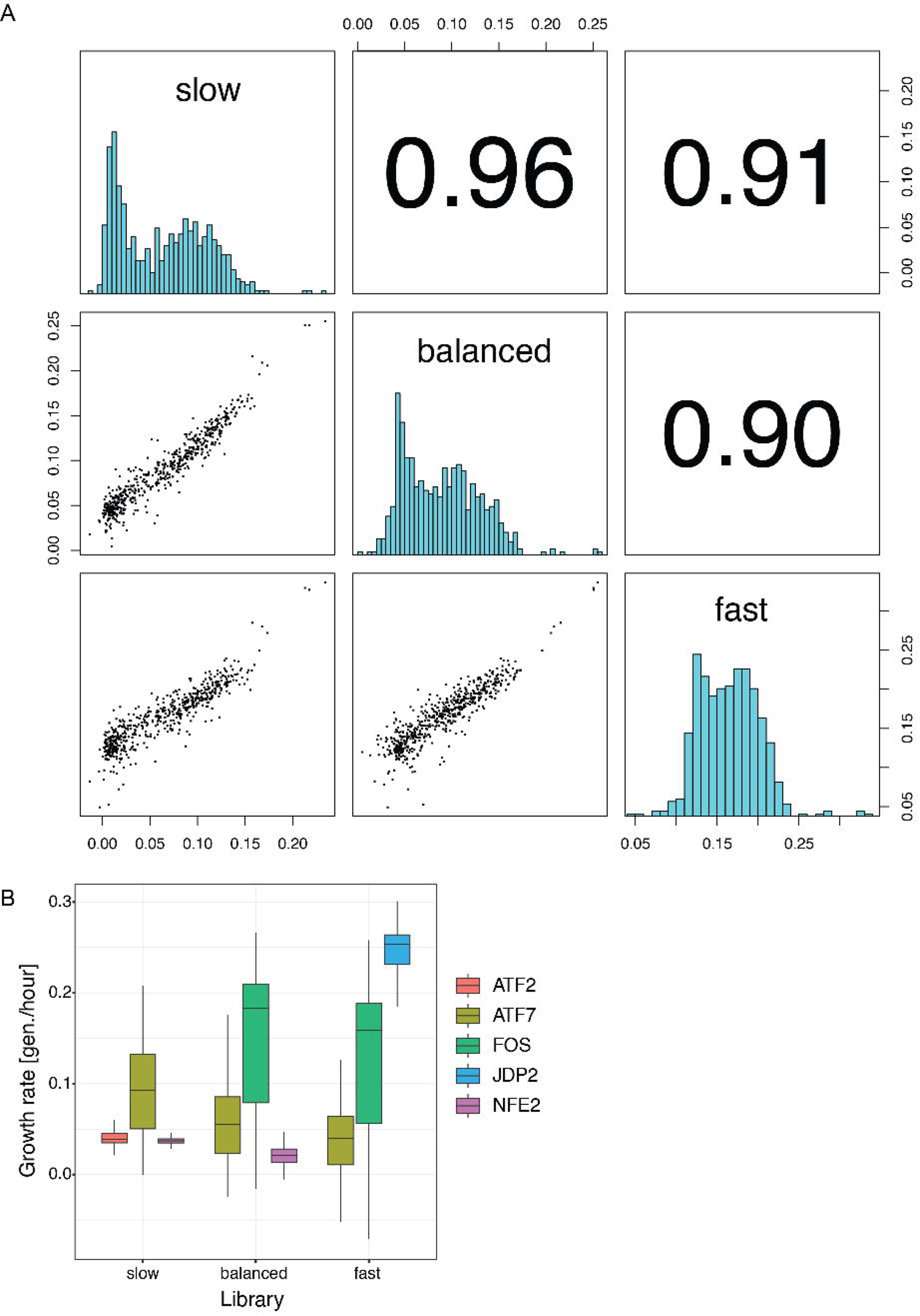
Growth rates are not independent of library composition. A. Pairwise correlations of overlapping components between growth rates obtained from deepPCAs performed with different libraries. Scatter plots are shown in the bottom left panels, histograms are shown on the diagonal, and Pearson correlation coefficients are shown in the upper right panels. B. Distributions of growth rates for individual components obtained from deepPCAs of slow-growing, balanced and fast-growing libraries.

## Conclusions

We have performed a multi-parameter optimization study of deepPCA to identify sources of non-linearity in the workflow. Parameters like amount of transformed DNA and timepoint of harvest were identified to introduce non-linearities in the data. While these can easily be accounted for by thermodynamic modeling, identifying these sources of non-linearity can help improve the modelling process. Adapting the experimental procedure to minimize this non-linearity would also help improve quantification accuracy.

We do note that all of these experiments were performed using rather small libraries. Although our findings should be generally applicable, it is essential to thoroughly plan every deepPCA beforehand and take into consideration potential additional sources of non-linearity that might come from different library sizes or properties of the proteins screened.

## Methods

### Yeast Strain

All experiments presented were performed in BY4742 (MATα *his3*Δ1 *leut2*Δ0 *lys2*Δ0 *ura3*Δ0).

### Media and buffer recipes

- LB: 25 g/L Luria-Broth-Base (Invitrogen, Waltham, MA, USA). Autoclaved 20 min at 121°C.
- LB-agar with 2X Ampicillin (100 μg/mL): 25 g/L Luria-Broth-Base (Invitrogen, Waltham, MA, USA), 7.5 g/L Agar, 1.2 g/L MgSO_4_ · H_2_O. Autoclaved 20 min at 121 °C. Cool-down to 45 °C. Addition of 100 mg/L Ampicillin.
- YPAD: 20 g/L Bacto-Peptone, 20 g/L Dextrose, 10 g/L Yeast extract, 25 mg/L Adenine. Filter-sterilized (Millipore Express ®PLUS 0.22 μm PES, Merck, Darmstadt, Germany).
- SC-ura: 6.7 g/L Yeast nitrogen base without amino acids, 20 g/L glucose, 0.77 g/L complete supplement mixture drop-out without uracil. Filter-sterilized (Millipore Express ®PLUS 0.22 μm PES, Merck, Darmstadt, Germany).
- SC-ura/ade/met: 6.7 g Yeast nitrogen base without amino acids and folic acid, 20 g/L glucose, 0.74 g/L complete supplement mixture drop-out without uracil, adenine and methionine. Filter-sterilized (Millipore Express ®PLUS 0.22 μm PES, Merck, Darmstadt, Germany).
- SORB: 1 M sorbitol, 100 mM LiOAc, 10 mM Tris-HCl pH 8.0, 1 mM EDTA pH 8.0. Filter-sterilized (Millipore Express ®PLUS 0.22 μm PES, Merck, Darmstadt, Germany).
- Plate mixture: 40 % PEG3350, 100 mM LiOAc, 10 mM Tris-HCL pH 8.0, 1 mM EDTA pH 8.0. Filter-sterilized (Millipore Express ®PLUS 0.22 μm PES, Merck, Darmstadt, Germany).
- Recovery medium: YPAD + 0.5 M sorbitol. Filter-sterilized (Millipore Express ®PLUS 0.22 μm PES, Merck, Darmstadt, Germany).
- Standard competition medium: SC-ura/ade/met + 200 μg/mL methotrexate (BioShop Canada Inc., Canada), 2 % DMSO.
- DTT buffer: 0.1 M EDTA-KOH pH7.5, 10 mM DTT
- Zymolyase buffer: 20 mM K-phoshpate pH 7.2, 1.2 M sorbitol, 0.4 mg/mL Zymolyase 20T (amsbio, USbiological), 100 μg/mL RNAse A
- Methotrexate (Sigma Aldrich/Merck, Darmstadt, Germany)

### Plasmid library construction

The libraries used in this study were derived from [11].

To study the effect of harvesting the cells after different growth generations, we used the exact same library as in [11]. For this, an intermediate library of all single amino acid variants of the JUN zipper region fused to the DHFR c-terminal half (referred to as FR-tag) and identifiable by 24 bp randomly generated molecular barcodes was created. Barcode and variants were associated as described in [11]. This library was combined with a pooled collection of plasmids carrying all 54 wt bZIPs fused to the DHFR N-terminal half (referred to as DH-tag) and identifiable by a Sanger sequencing-confirmed molecular barcode of 20 bp. The two intermediate libraries were cloned together using high-efficiency cloning and standard restriction enzyme-mediated cloning with enzymes AvrII (New England Biolabs, Ipswitch, MA) and HindIII-HF (New England Biolabs, Ipswitch, MA) as described in [11]. For all barcodes, see Supplementary Tables 11&12.

The other experiments were performed with a sub-library of the above one. A standard balanced library was created combining the full JUN variant prey library with 3 wt bZIPs as bait. These were chosen based on their wt interaction strengths with wt JUN that were derived from [11]. To achieve a library of balanced growth rate, the JUN library was combined with ATF7, a medium-strength interactor of JUN, FOS, a strong interactor of JUN, and NFE2, a weak interactor of JUN, as bait using the same cloning strategy as above. This library was called balanced JUN library.

For testing different library compositions, two additional libraries were created. For a slow-growing library, the JUN variant library was combined with ATF7, NFE2 and one additional weak interactor of JUN, ATF2, as bait. This library will be referred to as slow JUN library. For a fast-growing library, the JUN variant library was combined with ATF7, FOS and one additional strong interactor of JUN, JDP2, as bait. Again, the same cloning strategy as above and as described in detail in [11] was applied. This library will be referred to as fast JUN library.

### deepPCAs

#### Large-scale yeast transformation

Yeast cells of the strain BY4742 were grown to saturation overnight in YPAD. For individual replicates, single colonies were used to inoculate the culture. In the morning, different sizes of pre-culture were inoculated at an OD600 of 0.3 in 175 mL YPAD medium. The cells were grown for ca. 4 hours until they reached an OD600 between 1.2 and 1.4. (Because of the larger library size, all volumes at each step in the different generation times deepPCA were doubled. The exact protocol for transformation of this library can also be found in [11].) Next, the cells were harvested for 5 min at room temperature (RT) and 3,000 g. The pellets were washed first with 50 mL H2O and then 50 mL SORB and finally resuspended in 7 mL SORB and incubated on a wheel at RT for 30 min. To each sample, 175 ul of 10 mg/mL salmon sperm DNA (Agilent Technologies, Santa Clara, CA) were added, and the tubes were mixed well. Afterwards, DNA was added to the samples according to Table 1 followed by thorough mixing by shaking the tube.

**Table 1:** Overview of DNA amounts and libraries used in individual experiments.

| Experiment | Amount of DNA transformed per<br>175 mL per-culture | Library used |
| --- | --- | --- |
| measurement of<br>percentage of double<br>transformants | 250 ng, 500 ng, 750 ng, 1 µg<br>(details see below) | equimolar pool of similar<br>sized plasmid backbones<br>with URA3 and LEU2<br>auxotrophic selection<br>markers |
| transformation with<br>different amounts of<br>DNA deepPCA | 100 ng, 333 ng, 1 µg, 3 µg, 20 µg | balanced JUN library |
| lag/no-lag phase<br>deepPCA | 1 µg | balanced JUN library |
| different MTX<br>concentrations/brands,<br>fitness effect in DMSO<br>deepPCA | 1 µg | balanced JUN library |
| different generation<br>times deepPCA | 3.5 µg | full JUN vs. 54 wt bZIPs<br>library [11] |
| different library<br>compositions deepPCA | 1 µg | balanced JUN library, slow<br>JUN library, fast JUN library |

Afterwards, 35 mL of Plate Mixture were added to each sample and the samples were incubated at RT on a wheel for another 30 min. 3.5 mL of DMSO were added to each sample followed by a 20 min-heatshock in a 42 °C water bath. To ensure homogeneous distribution of the heat within the sample, the tubes are inverted a few times after 1, 2.5, 5, 7.5, 10 and 15 min. The cells were then spun down for 5 min at RT and 3,000 rpm. After thorough removal of the liquids, they were resuspended in recovery medium and incubated for 1 hour at 30 °C without agitation. The recovered cells were finally spun down for 5 minutes at RT and 3,000 g, the supernatant was removed, and the pellet was resuspended in 350 mL of SC-ura for selection 1. Additionally, cells were plated on SC – ura plates for counting the amount of transformants. For measuring the percentage of double transformants, cells were additionally plated of SC-leu and SC-ura-leu (see below). Selection 1 was grown for ca. 48 hours until the cells reached saturation.

### Measurement of percentage of double transformants

As plasmid with URA3 resistance, a plasmid derived from pRS416 [24] was used. As plasmid with LEU2 resistance, we used a lab-internal plasmid called pDL00266 that was derived from pRS415 [24] and was of similar size than pRS416 to facilitate equimolar mixing. An equimolar pool of the two plasmids was created and was transformed according to the protocol above. To measure the number of transformants and double transformants, different volumes of cells that would result in countable amounts of colonies on the plates were plated after transformation according to Table 2 and grown for 2 days at 30 °C. Additionally, selection 1 cultures of 350 mL were set up for all samples. Each sample was performed in duplicate.

**Table 2:** Plating scheme for counting of double transformants directly after transformation.

| Sample no. | DNA amount | Plating SC - URA and SC -<br>LEU | Plating on SC -URA/-<br>LEU |
| --- | --- | --- | --- |
| 1 | 250 ng | 2 x 40 ul, 2 x 110 ul | 2 x 400 ul, 2 x 1.1 mL |
| 2 | 500 ng | 2 x 35 ul, 2 x 100 ul | 2 x 350 uL, 2 x 1 mL |
| 3 | 750 ng | 2 x 20 ul, 2 x 60 ul | 2 x 200 ul, 2 x 600 ul |
| 4 | 1 ug | 2 x 10 ul, 2 x 50 ul | 2 x 100 ul, 2 x 500 ul |

After 2 days, the colonies on the plates were counted, the total amount of transformants per 350 mL selection culture was calculated and for each individual sample, the percentage of double transformants was calculated using equation 1.

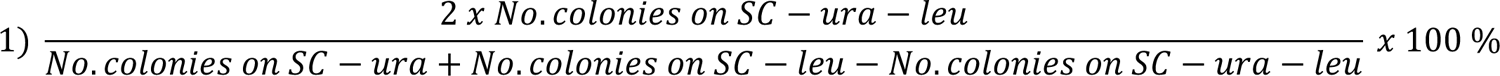

The averages of replicates and their standard deviations were calculated. The total number of transformants is represented by the denominator of equation 1 and corresponds to the sum of transformants on the -ura and -leu selections. Since these both also include double transformants that would hence be counted twice since present on the two plates, we then subtract the number of double transformants counted on the -ura/leu plates. The number of double transformants on the numerator of equation 1 is multiplied by two based on the assumption that 25% of double transformants received to ura plasmids, 25% received two leu plasmids and 50% received one of each, which are the only ones that can be counted on the - ura/leu plates.

Next, the saturated cultures of selection 1 were measured and the number of cells required to inoculate 200 mL of selection 2 in SC-ura-ade-met at OD600 of 0.1 was harvested. The cultures were inoculated and grown for ca. 12 hours until an OD of 1.2 was reached. Since there was no selection for the maintenance of the -leu plasmid alone and it was expected to only observe pRS415 plasmids in double transformants together with the pRS416-derived plasmid, the second round of plating was only done on SC-ura and SC-ura-leu according to Table 3. The cells were again incubated at 30 °C for 2 days after which the colonies were counted and the total number of transformants was calculated.

**Table 3:** Plating scheme for counting of double transformants after selection 2.

| Sample no. | DNA amount | Plating SC - URA | Plating on SC -URA/-LEU |
| --- | --- | --- | --- |
| 1 | 250 ng | 2 x 2 ul, 2 x 5 ul of 1/100 X diluted culture | 2 x 200 ul of undiluted culture, 2 x 20 ul of undiluted culture, 2 x 2 ul of undiluted culture |
| 2 | 500 ng | 2 x 2 ul, 2 x 5 ul of 1/100 X diluted culture | 2 x 200 ul of undiluted culture, 2 x 20 ul of undiluted culture, 2 x 2 ul of undiluted culture |
| 3 | 750 ng | 2 x 2 ul, 2 x 5 ul of<br>1/100 X diluted<br>culture | 2 x 200 ul of undiluted culture, 2 x 20 ul of<br>undiluted culture, 2 x 2 ul of undiluted<br>culture |
| 4 | 1 ug | 2 x 2 ul, 2 x 5 ul of<br>1/100 X diluted<br>culture | 2 x 200 ul of undiluted culture, 2 x 20 ul of<br>undiluted culture, 2 x 2 ul of undiluted<br>culture |

For colony numbers see Supplementary Table 1.

The percentage of double transformants was now calculated using equation 2.

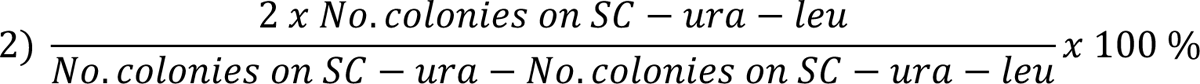

The averages of replicates and their standard deviations were calculated.

### Competition assay

All deepPCAs were started with selection 2: OD600 of the saturated cultures of selection 1 was measured and the number of cells required to inoculate 100 mL of SC-ura-ade-met at OD600 0.1 was harvested and re-suspended in the prepared medium. For the screen comparing the different harvesting timepoints that was performed with a larger library, 2 L of selection 2 were inoculated. The cells were grown at 30 °C and 200 rpm for ca. 12 hours until they reached an OD600 of 1.2. According to the condition tested, the competition assays were then performed slightly differently.

### Transformation with different amounts of DNA and different library compositions

100 mL of competition medium were inoculated at OD600 0.05 and grown at 30 °C and 200 rpm until they reached an OD600 of about 1.6. The remaining cells from selection 2 were harvested at RT or 4°C for 5 min at 3,000 OD, the pellet was washed twice with sterile H_2_O and then frozen at −20 °C. When the competition culture had reached the desired OD600, the cells were harvested in the same way and frozen.

### lag phase vs. no lag phase

For the + lag phase sample, 100 mL of competition medium were inoculated at OD600 0.05 and grown at 30 °C and 200 rpm until they reached an OD600 of about 1.6. The remaining cells from selection 2 were harvested at RT or 4°C for 5 min at 3,000 OD, the pellet was washed twice with sterile H_2_O and then frozen at −20 °C. For the - lag phase sample, the entire selection 2 culture was used to inoculate 100 mL of competition medium and was incubated at 30 °C and 200 rpm for 2.5 hours. From this pre-competition culture, another 100 mL of competition medium were inoculated at OD600 0.05 and grown at 30 °C and 200 rpm until they reached an OD600 of about 1.6. The remaining pre-competition culture was harvested as described above. All competition cultures that had reached the desired OD600 were harvested in the same way.

### Different MTX concentration/brands/DMSO

The assay was performed as described in “Transformation with different amounts of DNA” above, but the composition of competition medium was slightly adjusted depending on the sample (Table 4). To make sure that the same amount of DMSO would be added to each sample (2 mL), different stock concentrations of DMSO had been prepared beforehand.

**Table 4:** Concentration and brand of MTX used for different conditions in the deepCPA.

| Sample Name | Final concentration / brand MTX |
| --- | --- |
| DMSO | 0 µg/mL |
| 1/20 X MTX | 10 µg/mL, BioShop Canada Inc., Canada |
| 1/4 X MTX | 50 µg/mL, BioShop Canada Inc., Canada |
| 1/2 X MTX | 100 µg/mL, BioShop Canada Inc., Canada |
| 1 X MTX | 200 µg/mL, BioShop Canada Inc., Canada |
| 2 X MTX | 400 µg/mL, BioShop Canada Inc., Canada |
| 1 X MTX Sigma | 200 µg/mL, Sigma-Aldrich, Merck, Darmstadt, Germany |

### Different generation times

To harvest cells at low generation times and still get a sufficiently high number of cells for the subsequent DNA extraction and sequencing library preparation, the competition cultures were inoculated at different OD600s as listed in Table 5.

**Table 5:** Inoculation and harvest OD600 for different numbers of generations.

| Generations to grow until harvest | Inoculation OD600 | OD600 at harvest |
| --- | --- | --- |
| 2 | 0.2 | 0.8 |
| 3 | 0.1 | 0.8 |
| 4 | 0.05 | 0.8 |
| 4.5 | 0.05 | 1.2 |
| 5 | 0.05 | 1.6 |

2 L of competition medium were inoculated at the indicated OD600s from selection 2 culture and grown until they reached the desired OD600. They were then harvested as described above.

### DNA extraction

We recently introduced a novel technique to extract plasmid DNA from yeast cells that gives us about 10-times higher plasmid enrichment than the previous method as described in Diss & Lehner, 2018.

DNA from Input and Output samples was extracted by first spheroblasting the yeast cells followed by plasmid Mini- or Midiprep according to the culture size using Qiagen kits (QIAGEN, Hilden, Germany). To spheroblast the cells, pellets were thawed and incubated in 4 mL per 100 mL culture 0.1 M EDTA-KOH pH7.5, 10 mM DTT at 30 °C and 180 rpm shaking for 15 min. The cells were harvested at RT and 2,500 g for 5 min and re-suspended in 4 mL per 100 mL culture of 0 mM K-phosphate pH7.2, 1.2 M sorbitol, 0.4 mg/ml Zymolyase 20T (amsbio, USbiological) and 100 ug/mL RNase A. Cells were incubated at 30 °C and 180 rpm shaking until spheroblasting was complete after approximately 2 hours. Spheroblasts were collected at RT and 2,500 g for 5 min and then re-suspended in 1.6 mL per 100 mL culture homemade buffer P1 (according to the manufacturers protocol, QIAGEN, Hilden, Germany). 1.6 mL per 100 mL culture of homemade buffer P2 (according to the manufacturers protocol, QIAGEN, Hilden, Germany) was added. The samples were mixed well by inversion. Finally, 1.6 mL per 100 mL culture of pre-cooled, commercial buffer P3 (QIAGEN, Hilden, Germany) were added and the samples were again mixed well.

### Miniprep for cultures of 100 mL

The mixtures were transferred to 5 mL reaction tubes (Eppendorf, Hamburg, Germany) and spun at max. speed for 20 min in a tabletop centrifuge. The supernatant was recovered and plasmid DNA was purified following the standard QIAprep Spin Miniprep Kit (QIAGEN, Hilden, Germany) protocol and eluted in 50 μl EB buffer.

### Midiprep for cultures of 2L

The larger cultures were purified using a slightly adjusted Miniprep protocol as described in [11].

To quantify the amount of plasmid DNA in the extracts, qPCRs were performed using primers specific for the plasmid backbones (oGD241, oGD242. A plasmid of similar size to the library and known concentration was used to make a standard curve. The plasmid was pre-diluted to 0.4 ng/ul and a dilution series of 6 sequential 1/5 X dilution steps was performed in triplicate. qPCR was performed with the X SsoAdvanced Univeral SYBR Green Supermix (Bio-Rad Laboratories, Hercules, CA, USA) according to the manufacturers protocol.

### Sequencing library preparation

A single round of PCR was performed using NEBNext® Multiplex Oligos for Illumina®(New England Biolabs, Ipswitch, MA) that anneal to the Illumina primer binding sites already present in the barcode region of the plasmid. Depending on the sequencing run type and the expected reads per sample, an at lest 10-fold excess of plasmid molecules over expected reads was used as template for the PCRs. The libraries were amplified with Q5 polymerase (New England Biolabs, Ipswitch, MA). 50 μl reactions were set up with the required number of plasmid molecules, 1 X Q5 reaction buffer, 0.25 μM primer mix, 200 μM dNTPs, and 0.5 μl Q5 polymerase. The PCR was run between 14 and 20 cycles, depending on the PCR efficiency of the individual sample, with an annealing temperature of 63 °C and an extension at 72 °C for 30 seconds. Amplicons were gel-purifed from a 1 % Agarose gel using the QIAquick Gel Extraction Kit (QIAGEN, Düren, Germany). After determining the concentration with Qubit (Invitrogen, Waltham, MA, USA), samples were pooled at equimolar ratios and the pool was purified once more using AMPure XP magnetic beads (Beckman Coulter, Indianapolis, IN) according to the manufacturers protocol using a 1:1 ratio of beads and sample. As a final quality control, qPCR was performed on the pool to confirm the concentration measured with Qubit using the KAPA Library Quantification Standards with Primer (Roche Sequencing Solutions Inc, Pleasanton, CA), and the fragment distributions were checked on a Bioanalyzer (Agilent Technologies, Santa Clara, CA). The final library was submitted for sequencing on an Illumina sequencer (different sequencers were used, namely NovaSeq, HiSeq, NextSet, and MiSeq; all from Illumina, San Diego, CA). 50 bp single-end sequencing was performed in all cases to cover the barcode region. To increase sequencing, PhiX phage library was spiked into the library at varying amounts between 10 and 25 %.

### Data analysis

Raw read counts were processed and transformed into count tables using the mutscan R package [25]. Samples with an average Phred score below 20 and incorrect amplicon structure were discarded at this step. Afterwards, the sequences were matched with the barcode table and only perfect matches were kept. All barcodes belonging to the same variant were summed up and finally, all rows containing less than 10 reads in any Input or less than 1 in any Output were discarded. For each pair, generation times were calculated using equation 3.

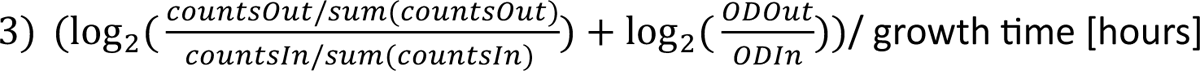

Figures were created in R Studio and Adobe Illustrator. All counts and average growth rates can be found in Supplementary Tables 2 to 10 as indicated in the main text.

## Availability of data and materials

### Data availability

Deep sequencing data is available at GEO with accession number GSE245485.

### Code availability

All custom scripts will be provided upon requests

## Acknowledgements

This work was supported by the Novartis Research Foundation and SNF Project grant 197593.

## Authors’ contributions

AMB and GD conceptualized the project. KS and AMB performed all the experiments with support from DK, KS and KKG. AMB analyzed the data. AMB and GD interpreted the results. AMB and GD wrote the manuscript.

## Declarations

### Ethics approval and consent to participate

Not applicable

### Consent for publication

Not applicable

### Competing interests

The authors declare no competing interests

---

**Supplementary Figure 1.**
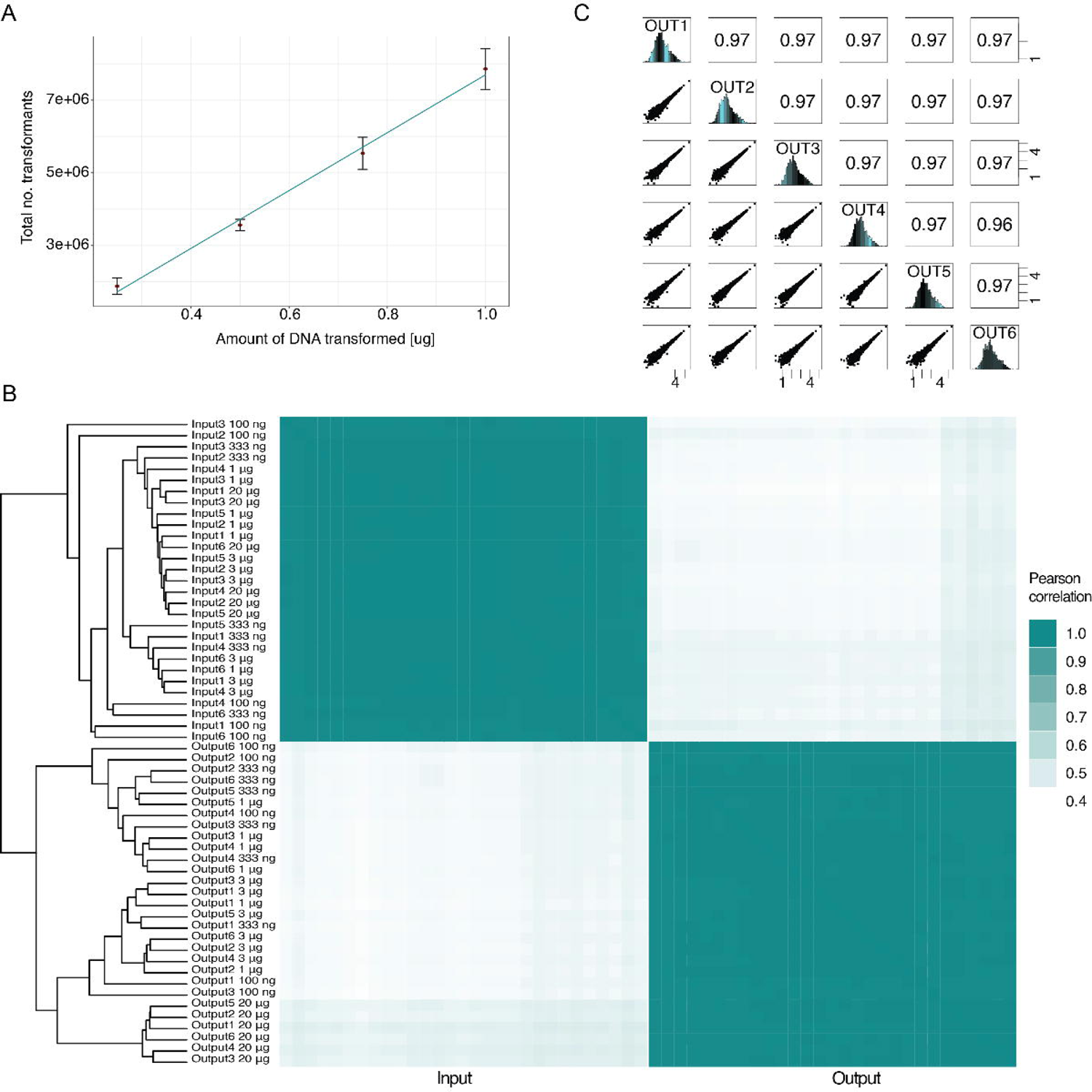
Count correlations between replicates transformed with different DNA amounts. A. Total number of transformants per transformation transformed with different amounts of DNA. B. Clustered correlation matrix between replicates of Input and Output samples from deepPCAs performed with cell that were transformed with different amounts of DNA. C. Pairwise correlations between counts of Output samples transformed with 20 μg of DNA. Scatter plots are shown in the bottom left panels, histograms are shown on the diagonal, and Pearson correlation coefficients are shown in the upper right panels.

**Supplementary Figure 2.**
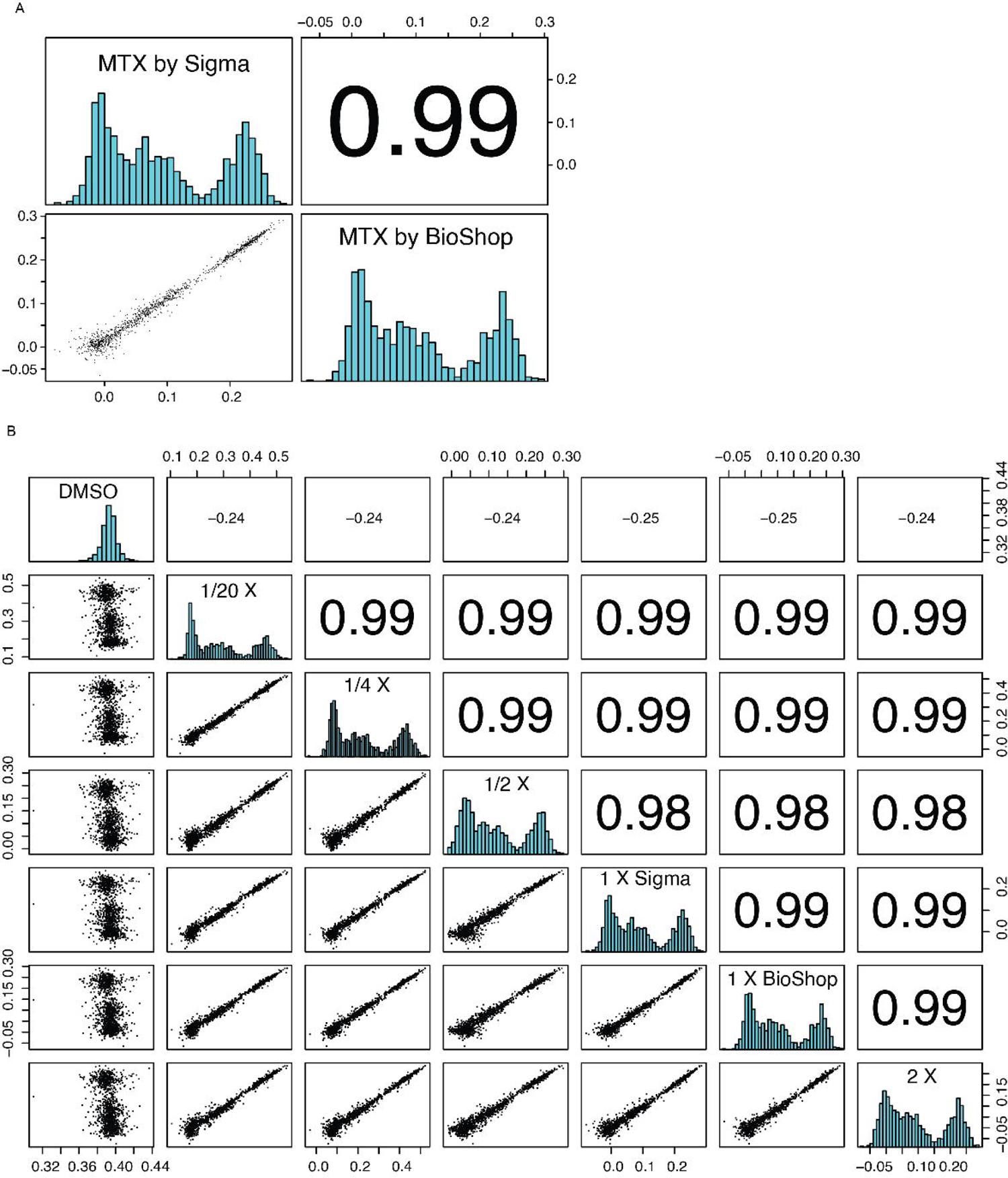
Correlation plots for experiments with different DNA amounts and brands. A. Pairwise correlation between growth rates obtained from deepPCAs performed with MTX from Sigma (Sigma Aldrich/Merck, Darmstadt, Germany) or BioShop (BioShop Canada Inc., Canada). The scatter plot is shown in the bottom left panel, histograms are shown on the diagonal, and the Pearson correlation coefficient is shown in the upper right panel. B. Pairwise correlations of growth rates between deepPCAs selected in DMSO, different concentrations and different brands of MTX. Scatter plots are shown in the bottom left panels, histograms are shown on the diagonal, and Pearson correlation coefficients are shown in the upper right panels.

**Supplementary Figure 3.**
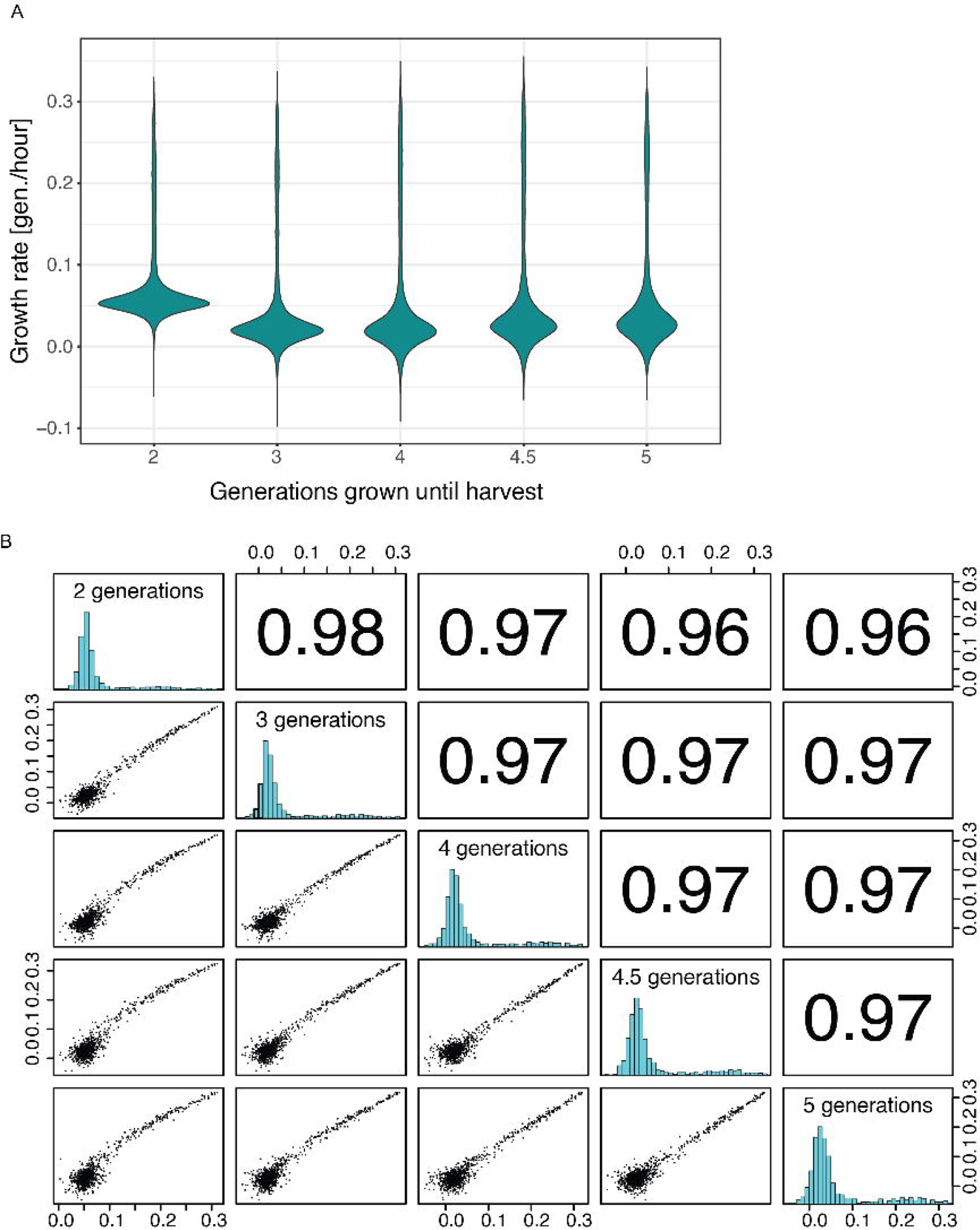
Harvest after different generations of growth affects growth rates non-linearly. A. Distribution of growth rates between deepPCA samples selected for different amounts of generations. B. Pairwise correlations of growth rates between deepPCA samples selected for different amounts of generations. Scatter plots are shown in the bottom left panels, histograms are shown on the diagonal, and Pearson correlation coefficients are shown in the upper right panels.

## Notes

### Competing Interest Statement

The authors have declared no competing interest.

